# High-resolution Ribosome Profiling Reveals Translational Selectivity for Transcripts in Bovine Preimplantation Embryo Development

**DOI:** 10.1101/2022.03.25.485883

**Authors:** Linkai Zhu, Tong Zhou, Rajan Iyyappan, Hao Ming, Yinjuan Wang, Michal Dvoran, Qi Chen, R. Michael Roberts, Andrej Susor, Zongliang Jiang

## Abstract

High resolution ribosome fractionation and low-input ribosome profiling of bovine oocytes and preimplantation embryos has enabled us to define the translational landscapes of early embryo development at an unprecedented level. We analyzed the transcriptome, polysome- and non-polysome-bound RNA profiles of bovine oocytes (GV and MII stage) and early embryos at 2-, 8-cell, morula, and blastocyst stage, and revealed four modes of translational selectivity: i. selective translation of non-abundant mRNAs, ii. active, but modest translation of a selection of highly expressed mRNAs, iii. translationally suppressed abundant to moderately abundant mRNAs, and iv. mRNAs associated specifically with monosomes. A strong translational selection of low abundance mRNAs encoding protein components involved in metabolic pathways and lysosome was found throughout bovine oocyte and preimplantation development. In particular, genes encoding components involved in mitochondrial function were prioritized for translation. Notably, transcripts encoding proteins regulating chromatin modifications selectively translated in oocytes. We found that the translational dynamics largely reflects transcriptional profiles in oocytes and 2-cell embryos, but observed marked shift in translational control in 8-cell embryos associated with the main phase of embryonic genome activation. Subsequently, transcription and translation become better synchronized in morulae and blastocysts. Together, these data reveal a unique spatiotemporal translational regulation that accompanies bovine preimplantation development.

**Significance Statement:** Translational control during preimplantation embryo development is poorly understood, mostly due to the scarcity of samples and the corresponding inability to analyze low quantities of these materials. By developing a low-input method, we have been able to explore the transcriptome, polysome- and non-polysome-bound RNA profiles of bovine oocytes (GV and MII stage) and preimplantation embryos at 2-, 8-cell, morula, and blastocyst stages. We reveal four different modes of translational selectivity, plus novel temporal regulatory mechanisms during early embryo development. The spatiotemporal translation dynamics of bovine oocytes and preimplantation embryos offer an entirely new insight into mammalian embryo development research and new possibilities for improving efficiency of assisted reproduction technologies (ARTs).

## Introduction

Preimplantation embryo development is a complex and precisely regulated process orchestrated by maternal stored mRNAs and newly synthesized transcripts that appear following embryonic genome activation (EGA) (1). In the last decade, transcriptome analyses of early mammalian embryos from multiple species have been comprehensively conducted and have established precise gene transcription programs during preimplantation development. However, the level of mRNA and the amount of its protein product often do not directly correlate (2), suggesting that the mRNAs detected from global transcriptomic profile do not necessarily represent their functional status in early embryo development. Although protein expression landscape of oocytes and preimplantation embryos has been characterized in mouse (3, 4), the proteomic analysis offers limited coverage and information due to scarcity of sample material available from oocytes and embryos and has not been explored in other mammalian species. More importantly, a central gap in our understanding of post-transcriptional regulation exists, namely how mRNAs are selected for spatially and temporal regulation during cell fate specification and in such as processes as oocyte maturation, fertilization, EGA and early differentiation. Thus, the understanding of mRNA translational dynamics may provide new insights of gene regulation during embryogenesis.

Accordingly, in some systems, ribosome profiling coupled to RNA-seq (Ribo-seq) has been developed to quantify ribosome occupancy and to analyze selective genome-wide mRNA translation (5, 6). However, the broad application of Ribo-seq has been slowed by its complexity and the difficulty of adapting it to low amounts of input material as encountered in mammalian oocytes and early embryos. Recently, two powerful single cell Ribo-seq assays have been developed. First, Ribo-STAMP (Surveying Targets by APOBEC (a cytosine deaminase that catalyzes RNA cytosine-to-uracil)-Mediated Profiling) uses APOBEC-mediated RNA editing to identify transcripts that have been associated with ribosomes (7). The other is the scRibo-seq that utilizes the micrococcal nuclease (MNase) to digest exposed RNA and capture the ribosome protected footprints (8). Both approaches require complex quality control and analysis due to the high “noise” observed with single cell data. In addition, it has been shown that mRNAs engaged in translation are bound by ribosomes, while dormant or stored transcripts are accumulated in diverse forms of ribonucleoprotein complexes and particles (9–11). It is also well known that actively translated mRNAs are bound by multiple ribosomes, the polysomes (12). The above-mentioned approaches limit analysis of the variation encountered in the different numbers of ribosome-bound mRNAs as a whole, while ignoring how the specific mRNAs are preferentially selected for translation. It should be noted that Zhang et al., have also optimized a low-input ribosome profiling (LiRibo-seq) approach and provided the first translational dynamics of mouse oocytes and preimplantation embryos (13), but again this study was confined to an analysis of ribosome bound mRNAs as a whole. On the other hand, an imaging-based approach performed on living Drosophila embryos has allowed the location and dynamics of translation of individual mRNA to be explored directly (14) and has opened up new avenues for understanding gene regulation during development; however, this technology is still in its infancy.

In our study, we have improved a recent advance of SSP-profiling (Scarce Sample Polysome Profiling) (15) based on physical polysome fractionation (5, 16). We substantially increased the resolution of the procedure to enable sequencing transcripts associated with monosomes and different sizes of polysomes extracted from bovine oocytes and preimplantation embryos. The data obtained has allowed us to study both genome-wide translational dynamics and translational selectivity mechanisms that accompany bovine early embryo development.

## Results

### mRNA translational landscapes in bovine oocytes and preimplantation embryos

Polysome profiling traditionally requires a large sample size in order to separate polysome-bound RNA and detect it by spectrophotometry, making the procedure challenging to apply to mammalian oocytes and embryos. SSP-profiling (fractionation of mRNAs based on the number of translating ribosomes by using sucrose-density gradients) overcomes some of these obstacles (15). The improved SSP-profiling followed by RNA-seq has allowed us to analyze mRNA translational profiles of bovine oocytes at the germinal vesicle (GV) and metaphase II (MII) stages, as well as preimplantation embryos at post-fertilization stages 2-, 8-cell, morula, and blastocyst stages (Figure 1A). For each, 200 oocytes or embryos were used, with the experiment performed twice. We collected ten equal volumes of fractions from the sucrose gradients for each developmental stage to provide a high resolution translatomic (transcripts associated with ribosomes from all fractionations) profile (Figure 1A). We conducted two analyses to validate the translatomic data. First, we assessed RNA isolated from each of ten fractions by qRT-PCR-based quantification of 18s and 28s ribosomal RNA (rRNA) (Figure S1), which allowed us to confirm the successful separation of free RNAs, 40s small ribosomal subunits, 60s large ribosomal subunits, monosomes (80s) and polysomes (*see methods*). The quantification of the 18s and 28s provided an assessment of the reproducibility of fraction collection (Figure S1). Second, Principal Component Analysis (PCA) and Pearson correlation analysis of translatomic data indicated consistent measurements between biological replicates in each fraction and across developmental stage (Figure 1B, Figure 2). Based on these analyses, we classified the ten fractionations into free RNAs (F1-F2), monosome bound (F3-F5 with F6-F7 as an intermediate stage), and polysome bound mRNA profiles (F8-10, regarded as polysome hereafter). In addition to ribosome profiling analysis, global transcriptome analysis was also performed on 20 oocytes (GV and MII stage) (n=3) and 20 embryos (n=3) at each developmental stage collected from the same batches used for ribosome fractionation and RNA-seq profiling. The transcriptomic data (triangles in Fig. 1C), especially in the PC1 dimension, appeared to organize roughly as two, seemingly distinct, groupings, namely the stages representing oocytes and 2-cell stage embryos and the stages from 8-cell to blastocyst (Figure 1C), which is consistent with the notion that bovine major embryonic genome activation (EGA) occurs at the 8-cell stage (17, 18).

**Figure 1.**
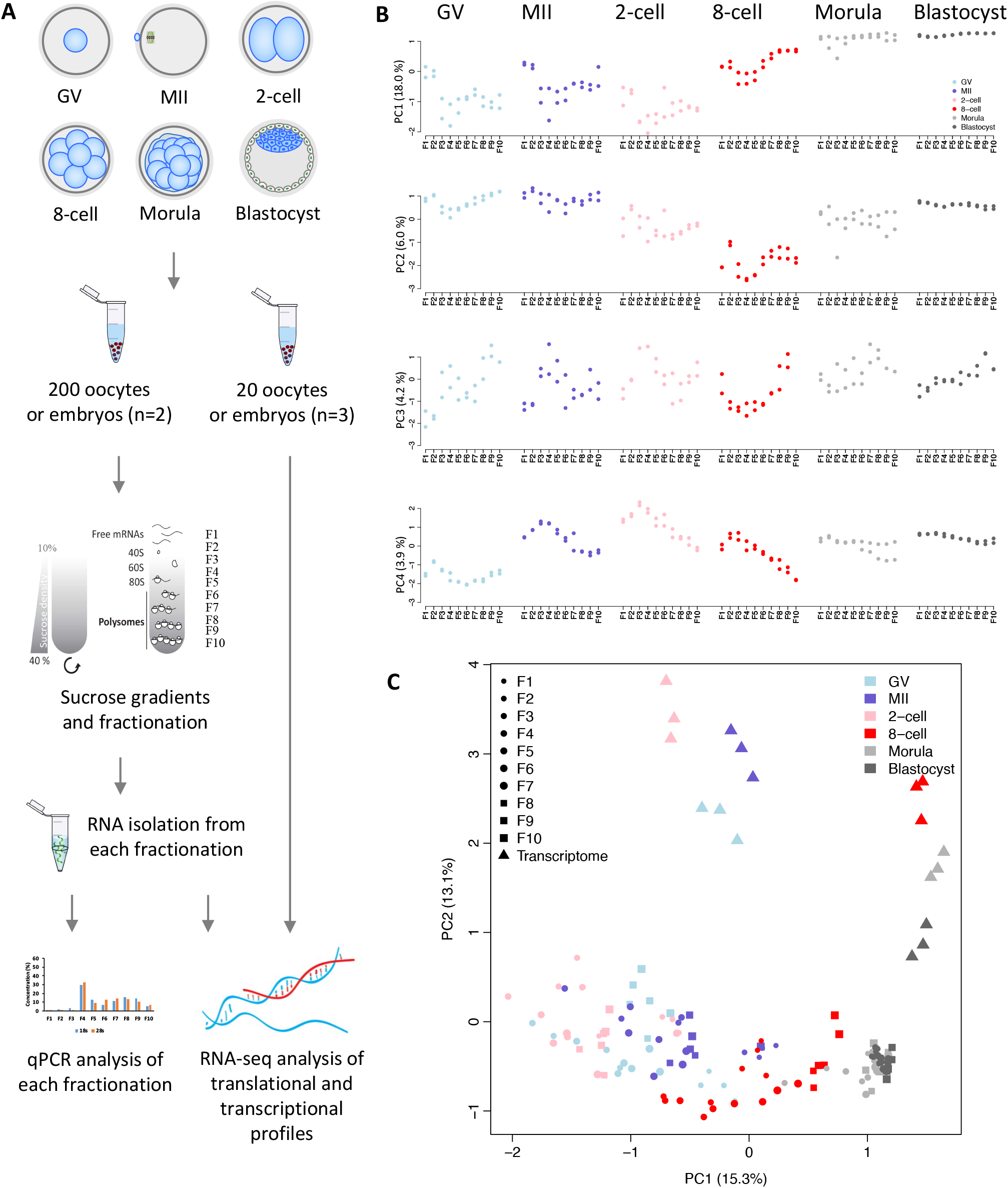
Genome-wide high-resolution ribosome profiling of bovine oocytes and early embryos. **A**. Scheme of genome-wide high-resolution polysome profiling in bovine oocytes and preimplantation embryos. **B**. Principal Component Analysis (PCA) of polysome- and nonpolysome-bound mRNA profiles in 10 fractions of bovine oocytes and early embryos. **C**. PCA analysis of translatomes (F1-F10) and transcriptomes of bovine oocytes and early embryos.

**Figure 2.**
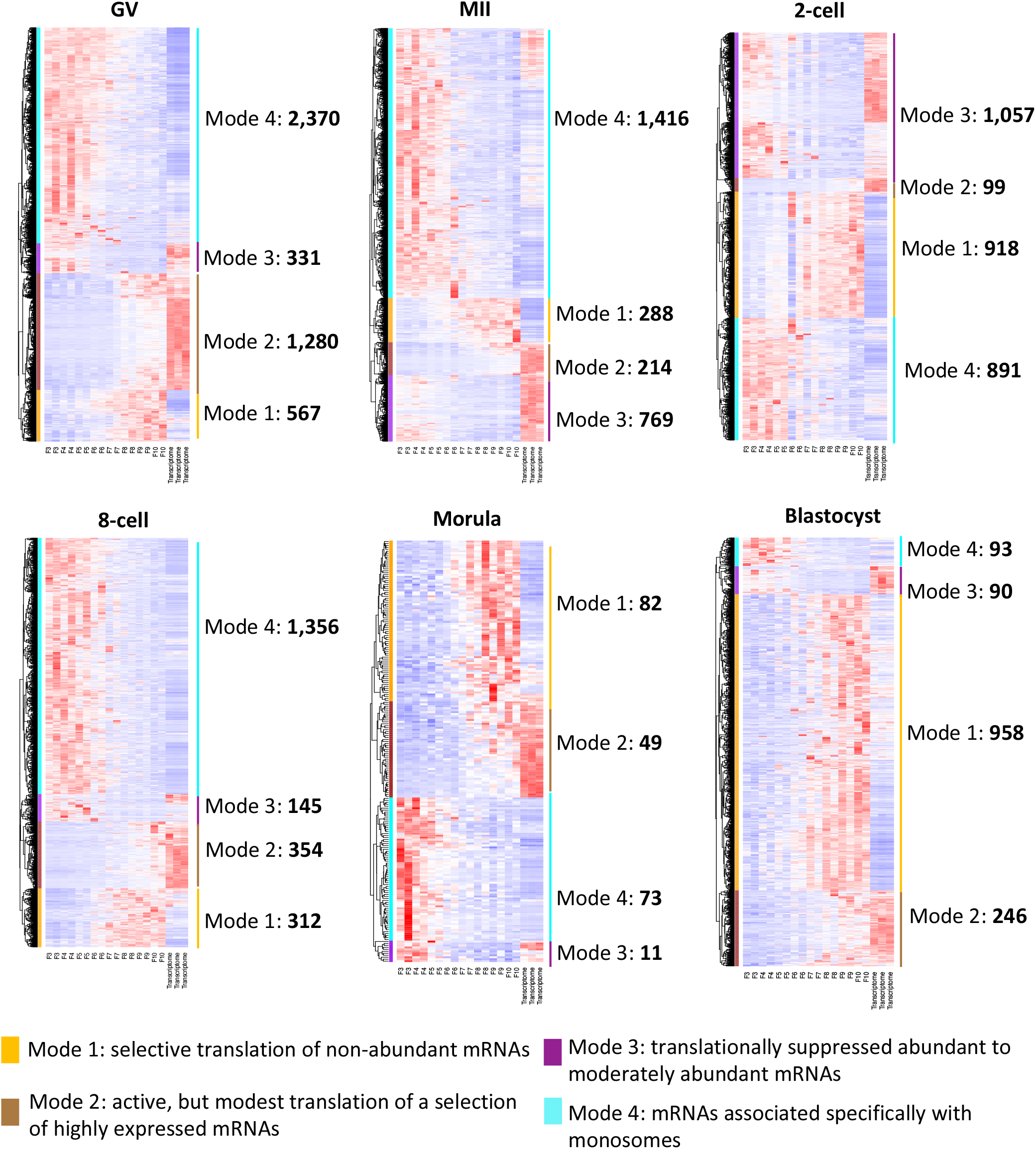
Diverse modes of translational selectivity during bovine oocyte and preimplantation development. Heat map showing (left panel) four modes of translational selectivity in bovine oocyte and preimplantation development. The color spectrum, ranging from red through white to blue, indicates high to low levels of gene expression. Mode 1. selective translation of non-abundant mRNAs (gold bar); Mode 2. active, but modest translation of a selection of highly expressed mRNAs (brown bar); Mode 3. translationally suppressed abundant to moderately abundant mRNAs (purple bar); and Mode 4. mRNAs associated specifically with monosomes (cyan bar). The numbers of genes identified in individual Modes of in each development stage are illustrated (right panel).

Overall, the translatome differed markedly from the transcriptome across different development stages (Figure 1C), suggesting discordance between the global transcriptome and actively translated mRNAs. Again, there was a separation of the data by stage. In particular the morula and blastocyst values were clustered together at the far right of the PC1 plot and well distanced from early-stage data which were clustered mainly towards the left of the PC1 plot and further separated from the rest of the developmental stages. Values for the 8-cell embryos fell somewhere in-between (Figure 1B, 1C). Our data also indicated that the changes in translatome appear to be gradual across fractions from F1 to F10 (Figure 1B, 1C), reflecting the continuous physical fractionation of mRNAs based on the number of translating ribosomes. While considerable differences existed between what transcripts are transcribed and what are translated, the various PCA analyses confirmed the largely similar trajectories of translatome and transcriptome dynamics during the development transition from oocytes to blastocysts, with a major shift occurring at the crucial 8-cell stage (Figure 1C).

### Diverse modes of translational selectivity during bovine oocyte and preimplantation development

To delineate the relationship between translation and transcription during bovine oocyte and preimplantation development, we assessed the correlation of all detected genes between translatome and transcriptome that had been generated from each of the six developmental stages. The F1-F2 fractions were excluded in order to allow us to focus on the translatome analysis (*see methods*). Overall, we found considerable differences between polysome-occupied (F8-10) and monosomaly-occupied mRNAs (F3-F5) over the course of development (Figure 2). We identified four modes of translational selectivity in each developmental stage: Mode 1: selective translation of non-abundant mRNAs (Figure 2, Gold bar); Mode 2: Active, but modest translation of a selection of highly expressed mRNAs (Figure 2, Brown bar); Mode 3: translationally suppressed abundant to moderately abundant mRNAs (Figure 2, Purple bar); and Mode 4: mRNAs associated specifically with monosomes (Figure 2, Cyan bar). A complete list of genes (Figure 2) from the four identified modes across bovine oocyte and preimplantation development are presented in Table S1, which should provide a valuable resource for others interested in translational regulation during bovine early embryo development.

Analysis of the functions of genes in Mode 2 (Active, but modest translation of a selection of highly expressed mRNAs) revealed a sequential progression of stage-specific gene networks. Gene enrichments shifted from cell division, chromosome organization, and mitotic nuclear division in oocyte (GV and MII), to embryonic cleavage and regulation of DNA replication in 2-cell, to translation in 8-cell, and finally to cell-cell adhesion and protein folding in morula and blastocyst when junctional complexes between cells become evident (Table 1).

**Table 1.**
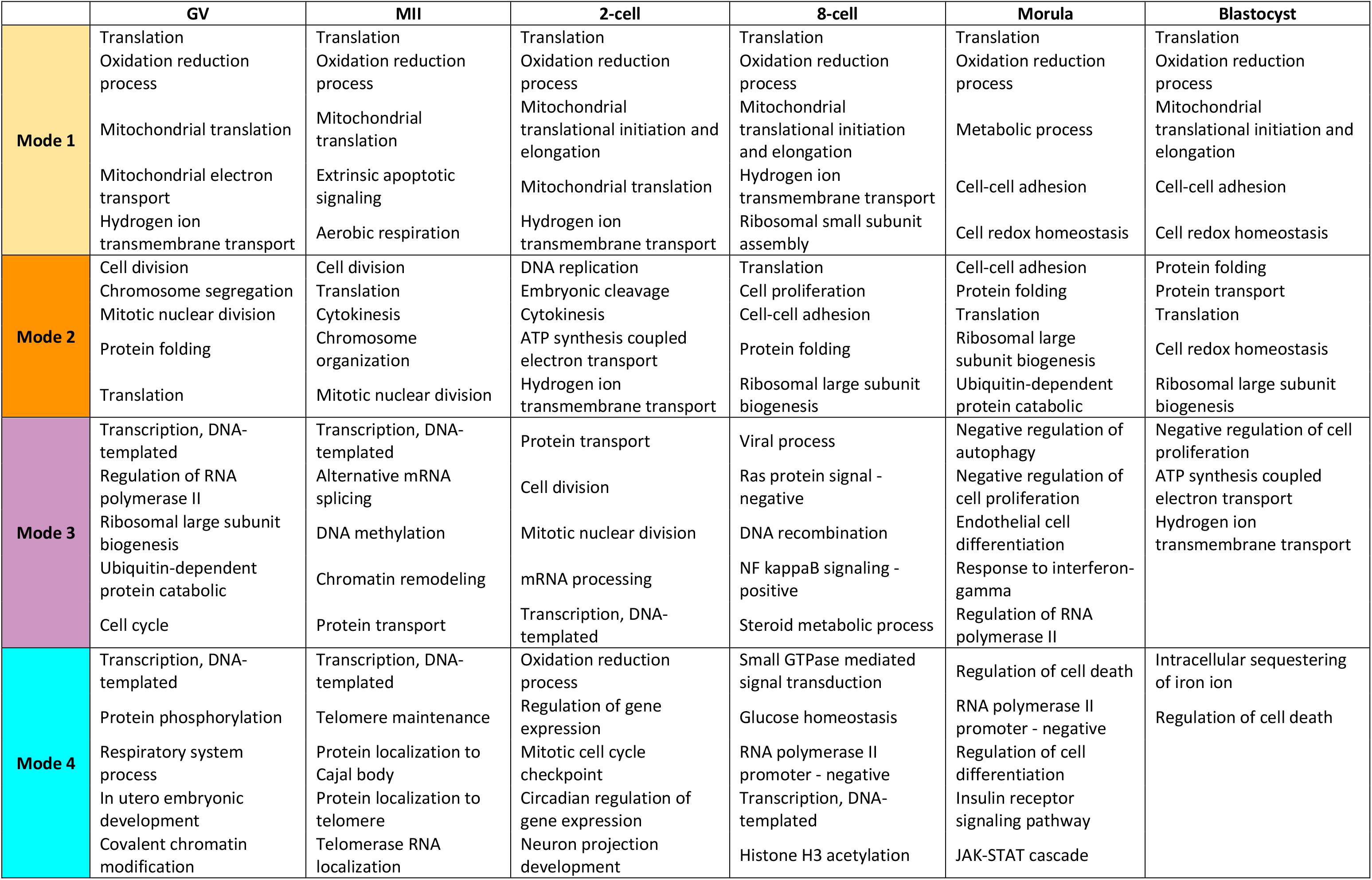
Top enriched GO terms associated with the genes from identified four modes of translational selectivity in each stage of bovine oocyte and preimplantation development.

Besides the gene groups that were highly expressed and actively translated, we identified second class of genes, sometimes relatively large in number, whose transcripts were of low abundance, yet that appeared to be actively translated, since they occupied polysomes (Mode 1: selective translation of non-abundant mRNAs; Figure 2). The common dominant biological processes represented in this mode included translation, oxidation-reduction process, mitochondrial translation; these functions were evident across all developmental stages (Table 1). Stage-specific programs included hydrogen ion transmembrane transport and apoptotic signaling at stages from oocyte to 8-cell and cell-cell adhesion and cell redox homeostasis at morula and blastocyst (Table 1).

The highly expressed but poorly translated transcripts (Mode 3: translationally suppressed abundant to moderately abundant mRNAs, Figure 2) were primarily involved in transcription, DNA-templated and RNA regulation in oocytes, protein transport and cell division at the 2-cell stage, viral process, and Ras protein signal transduction at the 8-cell stage, and negative regulation of autophagy and negative regulation of cell proliferation at morula and blastocyst (Table 1).

We also identified mRNAs occupying monosomes (Mode 4) from each developmental stage (Figure 2). Gene ontology analysis indicated significant gene enrichments relative to transcription, DNA-templated and protein phosphorylation at GV, transcription, DNA-templated and telomerase protein localization at MII, oxidation-reduction process and regulation of gene expression at 2-cell, small GTP signal transduction and glucose homeostasis at 8-cell, regulation of cell death and cell differentiation at morula, and finally intracellular sequestering of iron ion and regulation of cell death at blastocyst (Table 1).

In summary, our analysis captured four modes of translational selectivity for transcripts during bovine oocyte and preimplantation development. In particular, the analysis revealed gene activities (Modes 1 and 3) that could not be readily inferred from transcriptomic data alone.

### Translational control in bovine oocyte and preimplantation development

To gain insight into the broad translational regulation landscape across bovine oocyte and preimplantation development, we integrated the translatomes, i.e., transcripts associated with polysomes, with transcriptomes. The correlation between translatome and transcriptome was reasonably robust in GV and MII oocytes and in 2-cell embryos, but appeared strongest in GV oocytes where transcription is silenced, with the oocytes relying largely on abundant maternally stored RNAs, which are translated for oocyte growth and for the oocyte maturation process (Figure 3A). Translatomic data correlated less well with the transcriptome in MII oocytes than GV oocytes, when there remains a reliance on maternal transcripts but with more selective translation, possibly in preparation for fertilization (19). In contrast to the earlier stages, marked translational control is observed in 8-cell embryos (Figure 3A), indicating that polysome occupancy is poorly reflected in the transcriptome, most likely because this stage is when large scale transcription of the embryonic genome is being initiated, but the newly synthesized mRNAs may not yet fully occupy the ribosomal machinery. Of note, partial polysome-occupied mRNAs are being selected to translate immediately in the 8-cell embryo (Figure 3A), suggesting these genes are essential for the major EGA. Subsequent to the 8-cell stage, translation and transcription appear to become gradually better synchronized in morulae and particularly in blastocysts (Figure 3A), suggesting it is ready for the burst of protein production and cell proliferation that may be necessary to prepare the blastocyst for impending events, such as divergence of hypoblast and epiblast.

**Figure 3.**
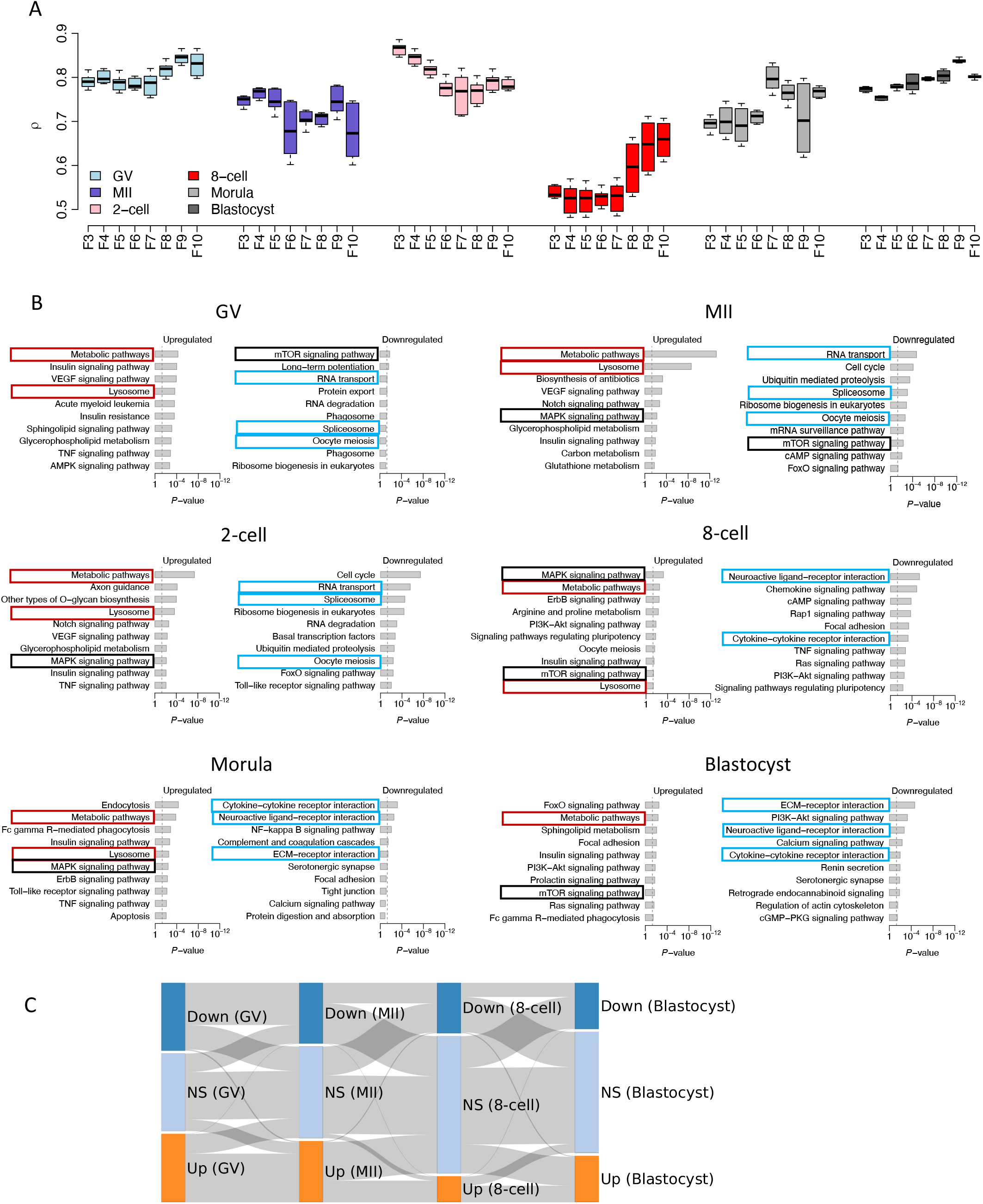
Translational control in bovine oocyte and preimplantation embryo development. **A**. Translational control is summarized by the correlation analysis of translatome (F3-F10) and transcriptome at each development stage. **B**. KEGG pathway analysis of the differentially expressed genes between polysome-occupied mRNAs (F8-F10) and transcriptome in bovine oocyte and preimplantation development. Red boxes highlight commonly pathways that are preferentially translated throughout bovine oocyte and preimplantation development; Cyan boxes highlight commonly downregulated pathways that are inactive translated before (GV, MII, 2-cell) or after (8-cell, morula, and blastocyst) major EGA stage; Black boxes highlight the most dynamic pathways that is translationally controlled throughout early development. Up- or down-regulated: FDR < 0.05, FC > 8; KEGG diseases pathways are excluded. **C**. Sankey diagram showing the up- and down-regulated genes (*FDR* < 0.05 and *FC* > 4) between polysome-occupied mRNAs (F8-F10) and transcriptome in each developmental stage. Down: down-regulated; Up: up-regulated; NS: not significant regulated.

To explore previously undefined translational dynamics in bovine oocyte and preimplantation development, we examined the pathways inferred from up- and down-regulated, polysome-associated transcripts compared to the transcriptome at each developmental stage using a stringent cutoff with false discovery rate [*FDR*] < 0.01 and fold change [*FC*] > 8 (Figure 3B). The broad term “metabolic pathways” and the narrower one “lysosome” were upregulated and, therefore, these mRNAs appear to be preferentially translated throughout bovine preimplantation development (Figure 3B). RNA transport, spliceosome and oocyte meiosis were pathways generally downregulated before the major EGA stage (GV, MII, and 2-cell), while commonly downregulated pathways at or after major EGA stage (8-cell, morula, and blastocyst) included various ligand-receptor interactions and extracellular matrix (ECM) receptor interaction (Figure 3B). Additionally, classical pathways, including those for mTOR and MAPK signaling, were the most dynamic pathways translationally controlled throughout early development (Figure 3B).

The data also revealed that the same polysome-occupied mRNAs in GV oocytes were largely retained in MII oocytes and only lost their translational selectivity at the 8-cell stage and beyond (Figure 3C), while the translationally suppressed mRNAs in GV oocytes were also essentially the same as the ones identified in MII oocytes and 8-cell stage embryos (Figure 3C).

### Translational switch occurs during bovine major EGA

To identify the genes with distinct translational trends as development progressed, we attempted to correlate the polysome-occupied mRNAs with stage. This analysis confirmed the dramatic translatome shift associated with the major EGA stage in the 8-cell embryo (Figure 4A, top panel). Until then, the up-regulated polysome-occupied transcripts detected in the later developmental stages, i.e., 8-cell, morula, and blastocyst, were significantly enriched for processes associated with translation, hydrogen ion transmembrane transport, cytoplasmic translation, ribosomal subunit assembly, and cell-cell adhesion (Figure 4A, bottom panel), while pathway analysis revealed a significant enrichment for ribosome assembly and oxidative phosphorylation (Figure 4A, bottom panel). The pathway analyses were also in agreement with these activities, especially in relation to energy metabolism. By contrast, the down-regulated polysome-occupied transcripts from the later stages, i.e., ones up-regulated in oocytes and 2-cell embryos, were associated with cell division, mitotic nuclear division, and DNA repair (Figure 4A, bottom panel), consistent with roles in oocyte maturation and early cleavage stages.The pathway analyses were also in agreement with these activities including cell cycle, RNA transport, oocyte meiosis, especially in relation to oocyte maturation (Figure 4A, bottom panel).

**Figure 4.**
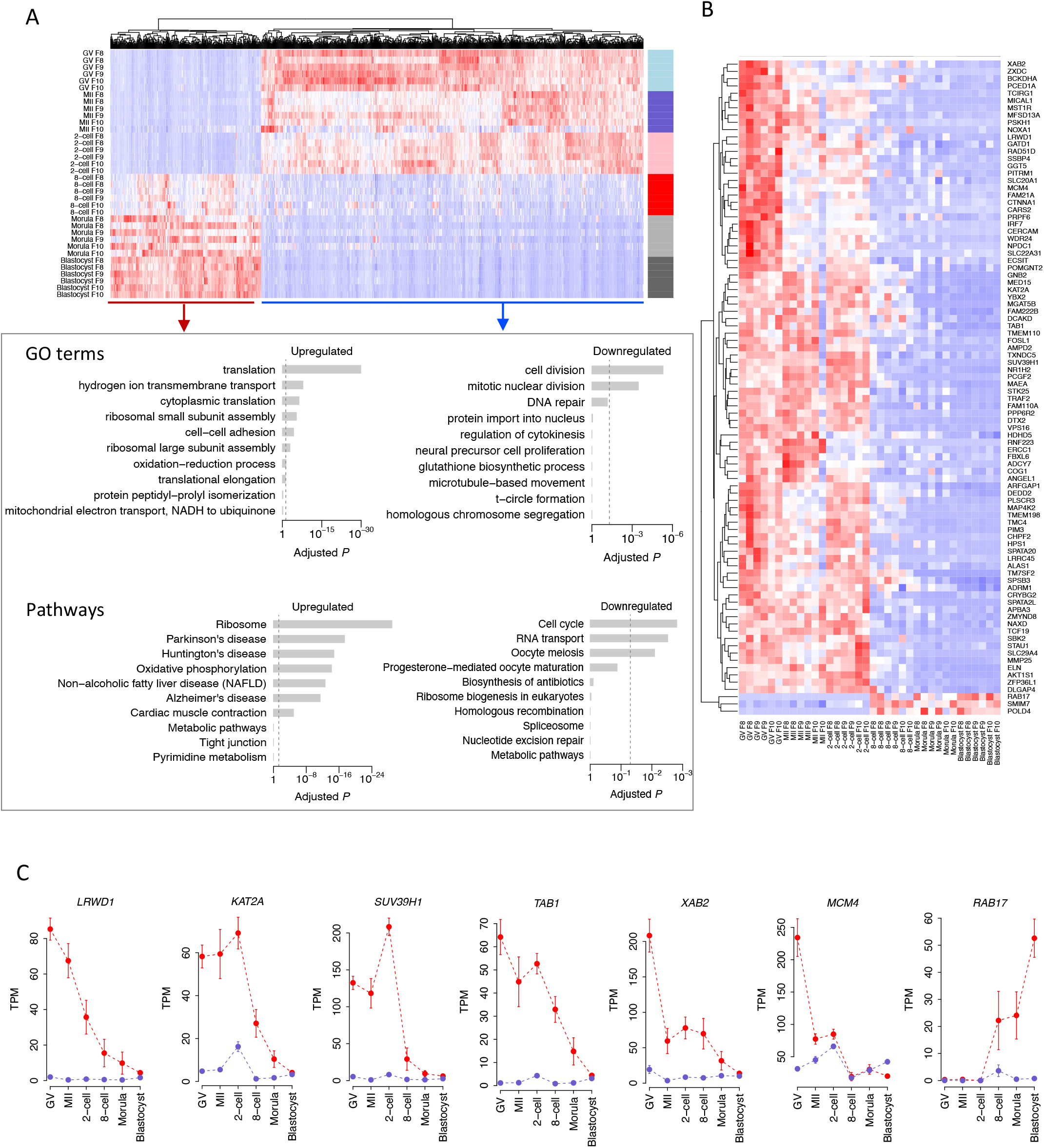
Translational switch occurs during bovine major EGA. **A**. Top panel: Heatmap showing the polysome-occupied mRNA (F8-F10) are correlated with developmental progression. The color spectrum, ranging from red through white to blue, represents high to low levels of gene expression. Bottom panel: Top enriched GO terms and KEGG pathways associated with up- (i.e., (up-regulation in 8-cell, morula, and blastocyst stages) or down-regulated (i.e., up-regulation in oocytes and 2-cell embryos) polysome-occupied genes towards to the developmental progression are presented. **B**. Heatmap of 90 prioritized genes with the most dynamic translational selectivity across bovine oocyte and preimplantation development. The color spectrum, ranging from red through white to blue, represents high to low levels of gene expression. **C**. Exemplary genes with distinct pattern between translation (Red line) and transcription (Blue line) in bovine oocyte and preimplantation development.

We then identified 90 genes that have the most dynamic translational selectivity across development (Figure 4B), of which most are actively translated in the oocyte to 2-cell stages and down-regulated thereafter. As examples, the top ranked down-regulated, polysome-occupied transcripts across developmental stages including *LRWD1, KAT2A, SUV39H1, TAB1, XAB2* and *MCM4* (Figure 4C). These genes all have their known functions linked to chromatin state. For example, *LRWD1* is a subunit of the origin recognition complex (ORC) and plays a role in heterochromatin organization and cell cycle control (20–23). *KAT2A* (also known as *GCN5*), a histone acetyltransferase, and *SUV39H1*, a histone methyltransferase that trimethylates lysine 9 of histone H3, are most highly studied histone enzymes and play pivotal roles in the epigenetic landscape and chromatin modification (24, 25). Given that a hallmark feature of a competent oocyte is chromatin condensation, the surprisingly highly selective translation of these genes in oocytes (both GV and MII) and the likely role of the translated proteins in maintaining the repressive heterochromatic state suggest that, in combination, these genes may have important function in the epigenetic control of bovine oocyte competence. Degradation of transcripts including *SUV39H1* and *TAB1* (Figure 4C) also have shown their essential roles in the maternal to zygotic transition (26, 27) and bovine preimplantation development (28–30), respectively. On the other hand, the top ranked up-regulated polysome-occupied transcripts across developmental stages is *RAB17* (Figure 4C), which belongs to a subfamily of small GTPases, plays an important role in the regulation of membrane trafficking (31). The translation of *RAB17*, which begins after the major EGA, is especially high at the blastocyst stage, when the trophectoderm lineage emerges and the blastocoel cavity forms. The other two transcripts with similar dynamics to *RAB17* are *SMIM7* and *POLD4* (Figure 4B), which encode a small integral membrane protein and a DNA polymerase subunit, respectively, appear to have nothing in common with each other or with *RAB17*. Their specific functions in bovine preimplantation development are unknown.

### Genes showing discordance between transcription and translation

We next analyzed the genes that showed contrasting trends in transcription *versus* translation (*FDR* < 0.05 and *FC* > 2) between stages during development from the oocyte to blastocyst (Figure 5A). Genes that had decreased transcription but an up-regulation of translation are represented by gold dots, whereas genes with increased transcription but decreased translation are in blue (Figure 5A). A total of 103 genes showed a decrease in transcript number, while at the same time had increased expression in the transition from GV oocyte to the MII stage (Table S2). Annotation of these genes revealed significant enrichment of mitochondrial translational initiation and translational elongation (Figure 5B). These findings, suggest that oocyte maturation requires a surge in the biosynthesis of mitochondrial components, which is consistent with the reported rise in aerobic metabolism accompanying oocyte maturation and gain of oocyte competence (32–34). By contrast, 65 genes had increased transcription but decreased translation (blue dots) during the 2-cell and 8-cell transition (Table S3). However conventional annotation analysis of these genes was not particularly informative (Figure 5B), although it must be assumed that some of these gene products play key roles in preparation for the major EGA occurring at the culmination of this transition.

**Figure 5.**
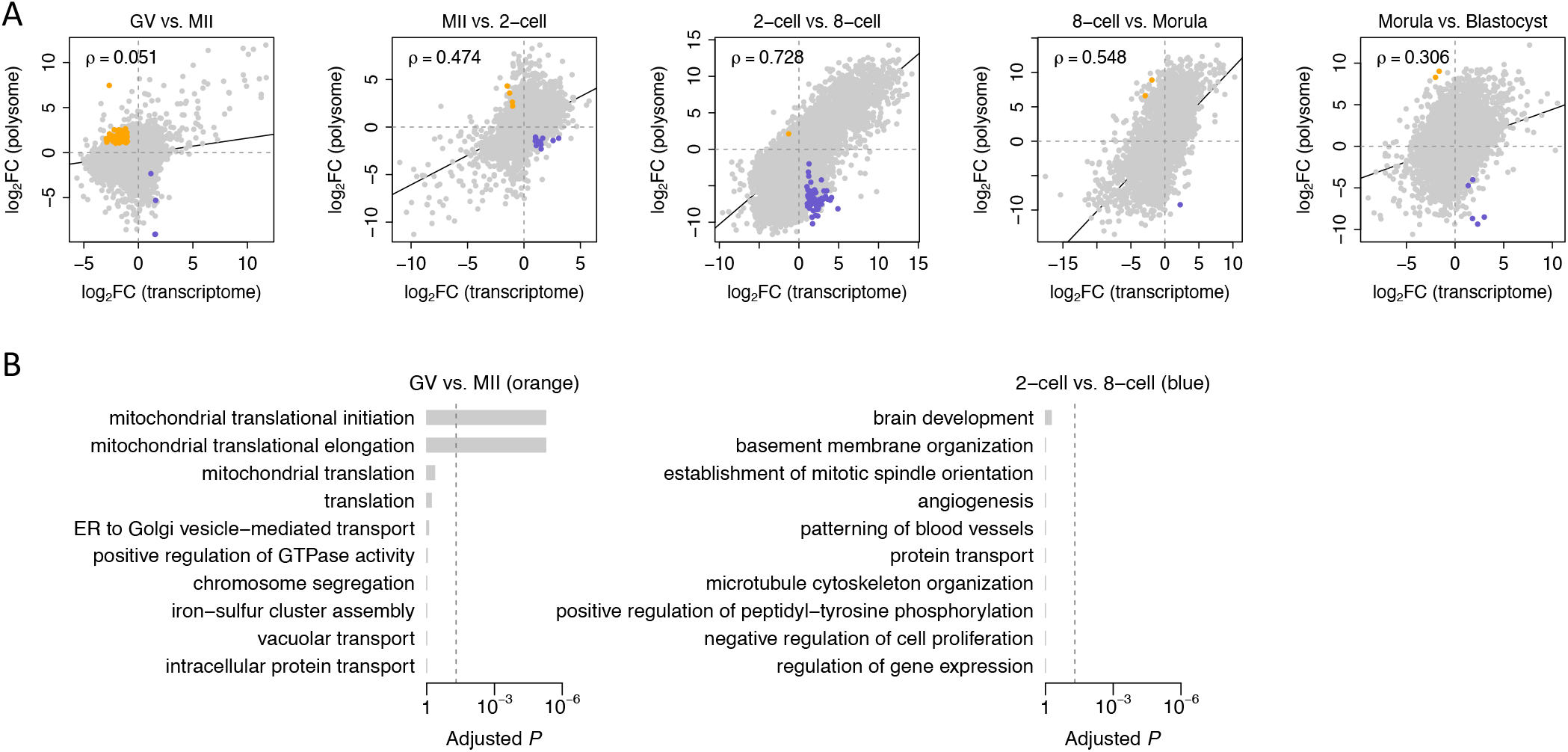
Genes showing discordance between transcription and translation. **A**. The **differential gene expression analysis** between polysome-occupied mRNAs and transcriptome in each developmental transition during bovine oocyte and preimplantation development. Gold dots represent genes that have decreased transcription but up-regulation of translations in each developmental transitions (*FDR* < 0.05 and *FC* > 2). Blue dots represent genes that have an increased transcription but decreased translation in each developmental transitions (*FDR* < 0.05 and *FC* > 2). **B**. The GO terms associated with the genes with decreased expression and up-regulated translations in MII compared to that of GV oocytes, and increased transcription but decreased translation during 2-cell and 8-cell transition, respectively.

We also identified the genes that are highly translated or transcribed at one particular stage but have low expression at other stages (Figure S2), suggesting that they likely have a specific regulatory function associated with a particular transition. The identification of these highly translated or transcribed genes at each developmental stage implies that each might have important regulatory roles during bovine embryonic programming. For example, well-known pluripotency genes *NANOG, KLF17, MYC*, and the interferon-response gene ISG15 are highly translated and transcribed at the 8-cell stage but much less so elsewhere. Again we observe the major EGA stage as particularly dynamic and one reflecting turbulent changes in gene expression.

## Discussion

Early embryonic loss greatly affects fertility of both humans (35, 36) and agriculturally important animals such as cattle (37), yet the underlying causes are, for the most part, unknown. A characterization of the molecular events accompanying the maturation of the oocyte, fertilization and the early cleavage stages of embryonic development may provide some insight into what can potentially go wrong in those pregnancies that fail in these early stages. Omics technologies have enabled in-depth analysis of molecular mechanisms of bovine preimplantation development including a catalog of the transcripts present (17, 18, 38–40) and the state of the epigenome (DNA methylation status (41, 42), chromatin dynamics (43, 44), histone modifications (45), and the range of small RNAs (46, 47)). However, the mRNA translation landscape and particularly the translational controls operating on specific mRNAs in oocytes and embryos remain largely unstudied. Here, we have developed a low input, high resolution, ribosome profiling approach and provided the genome-wide characterization of the important but often overlooked translational regulation process. The datasets, particularly when mined in further detail and integrated with epigenome information, are expected to greatly expand our understanding of the gene regulation mechanisms governing bovine embryonic development. Perhaps most importantly, significant discordance was frequently observed to exist between the linked processes of translation and transcription at each developmental stage of bovine early development, highlighting the importance of evaluating the translatome in addition to the more accessible transcriptome when studying development. Our study represents the first insights into mRNA translational dynamics and a comparison of the transcriptome with polysome- and non-polysome-bound mRNA profiles during mammalian oocyte and preimplantation development. In this regard, the bovine is recognized as a highly informative model for studying human embryo development (18, 43, 48, 49) on which such experiments are profoundly more difficult to conduct.

Our study was able to capture four diverse, although somewhat empirical, modes of translational selectivity for transcripts. In particular, Mode 1 (selective translation of non-abundant mRNAs) and Mode 3 (translational suppression of abundant to moderately abundant mRNAs) provides information that could not be inferred by transcriptome analysis alone. The mRNAs in Mode 1 provides a database for transcripts that are prioritized for translation relative to more abundant transcripts at each of the six stages of bovine preimplantation development examined. The identification of so many translationally suppressed, abundant to moderately abundant, transcribed genes, i.e., Mode 3 genes was somewhat surprising. The transcripts of these genes were largely absent from the polysome fractions, were most abundant in the oocyte and 2-cell stages, and diminished in number thereafter. A more detailed informatics analysis of these transcripts and an even a more comprehensive time course analysis seems warranted. One possibility is that their encoded proteins may be extremely stable or particularly efficient in their roles, so that low amounts of protein relative to mRNA are required for early development. Clearly any interpretation of the roles of the genes within either of these groups solely based on levels of their transcripts is bound to be incomplete. In conclusion, our study reveals unanticipated translational selectivity mechanisms operating on numerous genes across the genome. It identifies potentially important candidate regulators in embryonic programming that most likely have been overlooked in prior studies.

The analysis of genes in Mode 2 (active, but modest translation of a selection of highly expressed mRNAs), i.e., the ones that would likely predominate in a bulk transcriptomic analysis, revealed a sequential progression of stage-specific gene networks accompanying development. The data are largely consistent with the sequential changes revealed in our previous analysis of co-expressed genes in bovine oocyte and preimplantation embryo transcriptomes (18), but again reveal how transcriptomic data alone can be misleading and might overestimate the contribution of specific gene products to development. The transcripts that comprise Mode 4 contribute weakly to the transcriptome except at the MII oocyte stage (Figure 2), but appear to associate largely with monosomes and appear to be not actively translated at the stages examined. Perhaps this association provides a mechanism whereby excess transcripts can be temporarily laid aside but remain poised for future active translation. In other words, Mode 4 mRNAs associated specifically with monosomes may constitute a novel but temporary storage state for transcripts.

There is a major translational perturbance evident at the 8-cell stage of embryo development (Fig.1B, Fig. 3A, Fig. 4A). The data also show that there are consistent translational similarities between the GV oocyte, the MII oocyte, and the 2-cell stages (Figure 3A & C) but hat this pattern breaks down at the 8-cell stage, when the embryonic genome begins to contribute in a major way to the transcriptome. The transcripts identified in the early stages, i.e., GV oocyte to 2-cell embryo are presumed to be mainly, if not exclusively, of maternal origin and likely reflect similar physiological processes predominating throughout this developmental transition. Prior to the 8-cell stage and also subsequently at morula and blastocyst, translational dynamics are broadly correlated with the transcriptome. There is, however, a minor amount of transcriptional activity involving the embryonic genome at the 2-cell stage (17, 18) and this appears to correlate with high monosomes occupancy by mRNA (Fig. 1C; Fig. 3A). The implication of this observation is unclear.

Transcripts encoding proteins involved in mitochondrial function including oxidation reduction, electron transport chain, mitochondrial translational initiation and elongation, though not necessarily abundant are efficiently selected for translation at all stages of development (Table 1), reflecting the essential role of mitochondria in generating energy to support oocyte and embryo development (50). Transcripts encoding enzymes involved in a wide array of metabolic pathways are also preferentially translated at all stages, again not an unsurprising observation (51–57). Why these mRNA are so efficiently handled by the protein synthesis machinery remains unclear, however, but a deeper understanding of the metabolic networks operating during these stages might well facilitate improvement of media formulations for in vitro oocyte maturation and embryo culture and allow the development of biomarker assays for assessing oocyte and embryo competence.

It should be recognized that the oocytes and embryos used in this study are products of in vitro protocols. Neither oocyte maturation nor embryo development occur as efficiently under these conditions as they do in vivo, although new formulations are constantly being tested to improve the procedures. There is concern, therefore, that in vitro procedures not only contribute to some degree of developmental failure (58–60) but cause alterations in the transcriptome (61–63) and the translatome. Thus, the translational dynamic trajectory observed here in vitro might be somewhat different from that occurring in vivo. Nonetheless, in vitro fertilization and embryo in vitro culture are widely used in livestock species and in human in vitro fertilization programs. In particular, transfer of in vitro produced bovine embryos is a successful commercial practice in the cattle industry and has already surpassed the numbers of pregnancies achieved from in vivo-derived embryo transfers (www.iets.org). Therefore, the data obtained from the standard in vitro system used in the present paper has direct relevance to current practice in the clinic and on the farm. Although not currently feasible because of cost considerations relating to the numbers of oocytes and embryos required, a comprehensive comparison of translational dynamics of in vitro embryos with their in vivo counterparts might be of considerable interest.

Several new methods including Ribo-STAMP (7), LiRibo-seq (13), scRibo-seq (8), and imaging based SunTag (14) and RNA-puro-PLA (64), have recently opened avenues for understanding translational regulation with unprecedented cellular resolution. The main advantage of the SunTag, and RNA-puro-PLA, in particular, is to permit the localization and dynamics of mRNA translation to be observed at single molecule resolution. The development of the optimized SSP-profiling protocol described in the present study has enabled the characterization of translational status of mRNAs bound to different kinds of ribosomes (free subunits, monosomes, and polysomes) to be studied and has provided a more comprehensive pictures of translational control during bovine early development than ever achieved previously. Combined with highly sensitivity, high throughput mass spectrometry to permit full proteomics analyses (3, 65), our technology should be capable of providing detailed insights into the relative contributions of transcription, translation, and protein stability to the amounts of individual proteins in the developing embryo, as well as detailed regulatory mechanisms at play.

In summary, our study has revealed a previously unappreciated level of complexity in genome-wide translational selectivity mechanisms associated with oocyte maturation and embryo development. In particular the selective translation of non-abundant mRNAs for vital metabolic purposes throughout development, the stage-specific translational suppression of abundant to moderately abundant mRNAs, and the range of mRNAs associated with monosomes were particularly striking observations. Our work has filled a significant knowledge gap in the study of translational regulation over a period of rapid developmental change and provided an extensive data base that can be mined for more detailed insights into bovine oocyte and preimplantation development.

## Materials and Methods

### Bovine oocytes and *in vitro* embryo production

Germinal vesicle stage oocytes (GV oocytes) were collected as cumulus-oocyte complexes from follicles of 3-5 mm in diameter aspirated from slaughterhouse ovaries. BO-IVM medium (IVF Bioscience) was used for oocyte *in vitro* maturation. Maturation was conducted in four-well dishes for 22-23 hours at 38.5°C with 6% CO_2_ to collect MII oocytes. Cumulus cells were completely removed and maturation was confirmed by light microscopy examination. Cryopreserved semen from a Holstein bull with proven fertility was diluted with BO-SemenPrep medium (IVF Bioscience) and added to drops containing COCs with a final concentration of 2 × 10^6^ spermatozoa/ml. Gametes were co-incubated in 6% CO_2_ in air at 38.5°C for 18 hours. Embryos were then washed and cultured in BO-IVC medium (IVF Bioscience) at 38.5°C with 6% CO_2_. Different developmental stage embryos (2-cell, 8-cell, morula, and blastocyst) were then evaluated under light microscopy and only Grade 1 embryos by standards of the International Embryo Technology Society were selected for further study. Prior to oocyte and embryo collection, 100 ug/ml of cycloheximide (Sigma-Aldrich) was added into the culture for 10 minutes to stabilize and halt ribosomes on transcripts. Oocytes and embryos were then washed with D-PBS containing 1 mg/ml polyvinylpyrrolidone (PBS-PVP) and transferred into 50 μl droplets of 0.1% protease (Qiagen) to remove the zona pellucida. Oocytes and embryos were rinsed three times in PBS-PVP and confirmed to be free of contaminating cells, and then snap frozen in minimal medium and stored at −80°C until polysome fractionation.

### Isolation of ribosome-bound mRNA

Approximately 200 oocytes (GV or MII oocyte) or embryos at different developmental stages (2-, 8-cell, morula, and blastocyst) were combined with lysis buffer containing 10 mM HEPES (pH 7.5), 5 mM KCL, 5 mM MgCL2, 2 mM DTT, 1% Triton X-100, 100 μg/ml cycloheximide, complete EDTA-free protease inhibitor (Roche), and 40 U/ml RNase inhibitor. Oocytes and embryos were disrupted by zirconium silica beads (Sigma) in the mixer mill apparatus MM301 (shake frequency 30, total time 45s, Retsch). Lysates were cleaned by centrifugation in 10,000 x g for 5 min at 4°C and the supernatants were loaded into 10-40 % linear sucrose gradients containing 10 mM HEPES (pH7.5), 100 mM KCL, 5 mM MgCl_2_, 2mM DTT, 100 μg/ml cycloheximide, complete EDTA-free protease inhibitor, and 5 U/ml RNase inhibitor. Ultracentrifugation was carried out with a SW55Ti rotor and Optima L-90 Ultracentrifuge (Beckman Coulter). Ribosome profiles were recorded by ISCO UV absorbance reader. The overall quality of particular ribosome fractionation experiment was monitored by analysis of a parallel Hek293 cells sample. Ten equal fractions were then recovered and subjected to RNA isolation by Trizol reagent (Sigma).

### RT-PCR analysis

The RNA profile from each fraction was tested by qPCR analysis with 18s and 28s rRNA-specific primers to reconstruct a distribution of non-polysomal and polysomal RNA complexes in each profile (15). Briefly, 2 uL of RNA from each fraction were reverse-transcribed using 20U of M-MuLV Reverse Transcriptase (Thermo Scientific) and 0.3 ug of random hexamer primers in a reaction volume of 20 uL. cDNA synthesis was performed at 25 °C for 10 min and then in 37 °C for 5 min followed by incubation at 42 °C for 1 h and subsequent inactivation at 70 °C for 10 min. qRT-PCR experiments were performed in a LightCycler480® (Roche, Basel, Switzerland) and LightCycler480® SYBR Green I Master mix (Roche). The 10 uL reactions were performed in triplicate. Each reaction contained 2 uL of cDNA and 500 nM gene-specific primers (list of used primers are provided in Table S4). The amplification protocol was 95 °C for 5 min; 44 cycles of 95 °C for 10 s, 58 °C for 15 s, 72 °C for 15 s; followed by melting curve determination. For absolute qRT-PCR quantification, we created recombinant pCRTM4-TopoTM plasmids (Invitrogen, Carlsbad, CA, USA) containing 18S and 28S ribosome RNA PCR amplicons. The relative quantification mode was applied and the mean of 18s and 28s RNA level was used for normalization of each fractionation (Figure S1).

As described above, the RNA was separated in a sucrose gradient solution based on the number of ribosomes bound to the RNA. The 18s and 28s ribosomal subunits are central components of the 40s and 60s ribosomal subunits, respectively. Fractions 1 and 2 contained primarily free RNA; as a result, the concentration of the 18s and 28s would be expected to be low in comparison to the other fractions. Then, based on density, we anticipated high 18s rRNA and low 28s rRNA in fractions with 40s small ribosomal subunits, low 18s rRNA and high 28s rRNA in fractions with the 60s large subunits. Both would be present in the 80s monosomes and in polysomes, whose sizes would be evident from their alternating increasing content of both rRNAs. Therefore, the quantification of the 18s and 28s rRNA provides direct information on the reliability of fraction collection (15).

### Library preparation and RNA sequencing (RNA-seq)

The RNA-seq libraries were generated from individual fractions by using the Smart-seq2 v4 kit with minor modification from manufacturer’s instructions. Briefly, individual cells were lysed, and mRNA was captured and amplified with the Smart-seq2 v4 kit (Clontech). After AMPure XP beads purification, amplified RNAs were quality checked by using Agilent High Sensitivity D5000 kit (Agilent Technologies). High-quality amplified RNAs were subject to library preparation (Nextera XT DNA Library Preparation Kit; Illumina) and multiplexed by Nextera XT Indexes (Illumina). The concentration of sequencing libraries was determined by using Qubit dsDNA HS Assay Kit (Life Technologies) and KAPA Library Quantification Kits (KAPA Biosystems). The size of sequencing libraries was determined by means of High Sensitivity D5000 Assay in at Tapestation 4200 system (Agilent). Pooled indexed libraries were then sequenced on the Illumina HiSeq X platform with 150-bp paired-end reads.

A pool of 20 oocytes or preimplantation embryos (n=3) selected from the same batch in each developmental stage used for ribosome profiling were used to profile transcriptomes by RNA-seq following Smart-seq2 protocol as above described.

In total, we sequenced 138 RNA-seq libraries (120 ribosome bound mRNA libraries and 18 whole transcriptomes) and we generated approximately 40 million 150bp paired-end reads per sample. The raw FASTQ files and normalized read accounts per gene are available at Gene Expression Omnibus (GEO) (https://www.ncbi.nlm.nih.gov/geo/) under the accession number GSE196484.

### RNA-seq data analysis

The *Salmon* tool (66) was applied to quantify the genome-wide gene expression profile from the raw sequencing data, by using the Ensembl bovine genome annotation (ARS-UCD1.2). Transcript per million reads (TPM) was used as the unit of gene expression. The *edgeR* tool (67) was applied to identify differentially expressed genes. The *TMM* algorithm implemented in the edgeR package was used to perform normalization of the read counts and estimation of the effective library sizes. Differential expression analysis was performed by the likelihood ratio test implemented in the *edgeR* package.

In this study, the fractions of free RNAs (F1 and F2) were excluded because of the discontinuity with the other fractions in the global expression pattern (Figure 1B) and also because no ribosome bound RNA by qPCR analysis as described above. We anticipated the largely free RNA (not attached to any ribosome or protein) in F1-F2s might include microRNAs or non-coding RNAs, which play a significant function in early development based on recently studies (68–71). The inadequate annotation of such RNAs in bovine genome also limited the comprehensive characterizations in this study.

To understand the translational selectivity in each developmental stage, *Spearman*’s rank correlation test was applied to compute the relationship between gene expression and consecutive ribosomal fractions (F3-F10). The genes with significant gradual increase/decrease in expression and at least 2-fold change in expression between the upper (F8-F10) and lower (F3-F5) fractions were retained for further analysis.

All the conventional statistical analyses were performed based on the *R* platform. The “cor.test” function was used to perform *Spearman*’s rank correlation test. A linear model controlling for fractionation was applied to prioritize the polysome occupied genes with gradual increase/decrease in expression across the developmental stages using the “lm” function. If multiple testing should be accounted for, the “p.adjust” function was applied for *P*-value correction. Principal component analysis on the genome-wide gene expression profile was performed by using the “dudi.pca” function within the package “ade4”. All the heatmaps were plotted by the “heatmap.2” function within the package “gplots”. The gene ontology and pathway analysis were performed by means of the David tool (72).

## Acknowledgements and funding sources

This work was supported by the NIH Eunice Kennedy Shriver National Institute of Child Health and Human Development (R01HD102533) and USDA National Institute of Food and Agriculture (2019-67016-29863). A.S. is supported by the grant GACR 22-27301S.

## Supplementary Figure Legends and Tables

**Figure S1**. qRT-PCR analysis of the distribution of 18S and 28S rRNA in each fraction of bovine oocyte and preimplantation development.

**Figure S2**. Venn diagram showing the genes that are specifically and highly translated or transcribed in one particular stage across bovine oocyte and preimplantation development. The highly translation of genes (blue color) and the highly translation and most abundant genes (red color) specific to each development stage are listed.

**Supplementary table 1**. A complete list of genes from identified four modes of translational selectivity across bovine oocyte and preimplantation development. In each of developmental stage, spreadsheet lists genes from four different modes as specific by 1, 2, 3, and 4, respectively.

**Supplementary table 2**. A list of 103 genes that are identified to be down-regulated their expression, while up-regulated their actual translations in MII compared to GV oocytes (*FDR* <0.05 and *FC* > 2).

**Supplementary table 3**. A list of 65 genes that are identified to be up-regulated their expression, while down-regulated their actual translations in during 2- to 8-cell transition (*FDR* < 0.05 and *FC* > 2).

**Supplementary table 4**. Primers for qPCR analysis of 18s and 28s rRNA in the RNA from each fraction of bovine oocyte and preimplantation development.

## Notes

### Competing Interest Statement

The authors have declared no competing interest.

