## Supplementary figures and images for "High-resolution Ribosome Profiling Reveals Translational Selectivity for Transcripts in Bovine Preimplantation Embryo Development"

### Figure S1

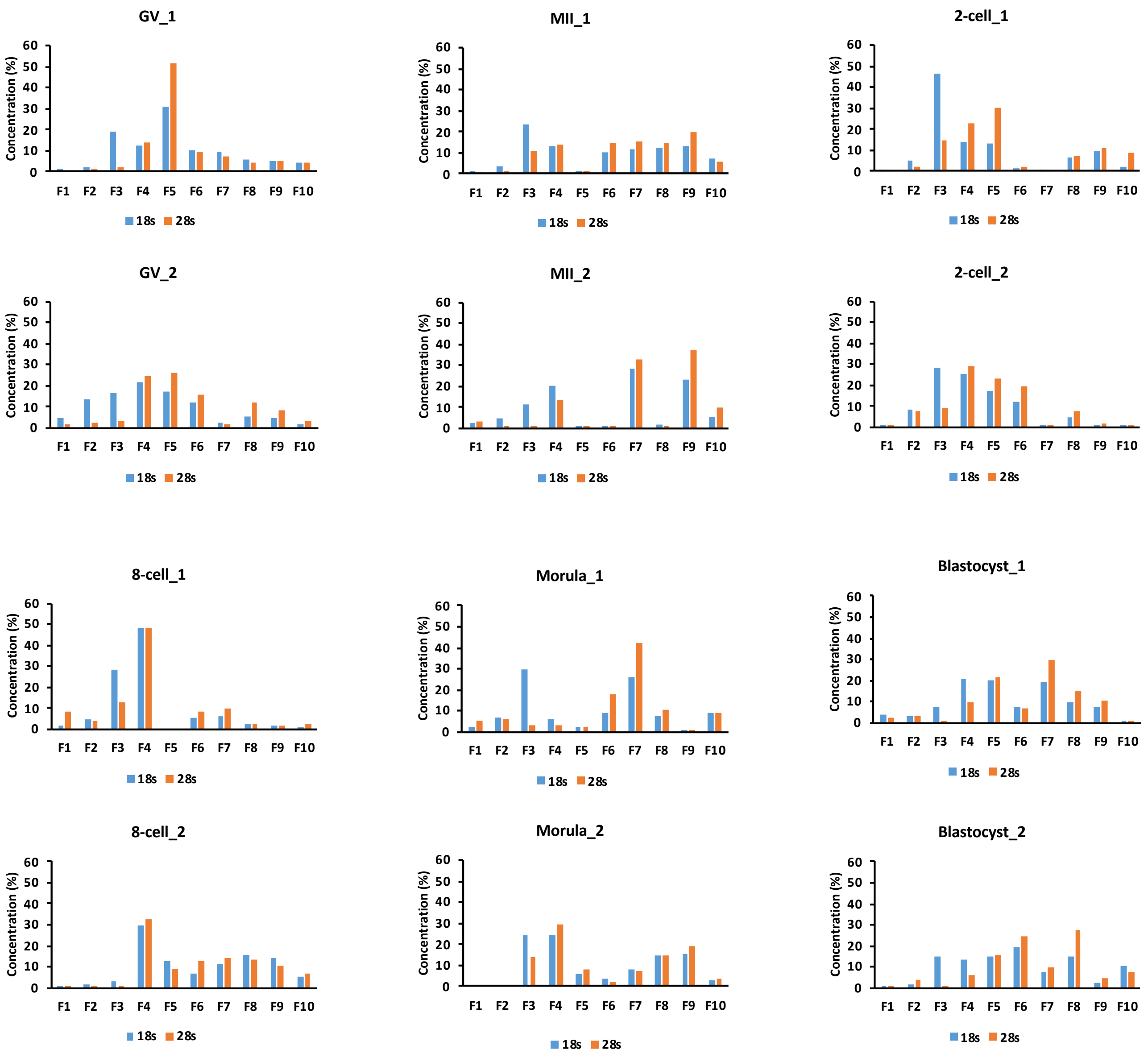

### Figure S2

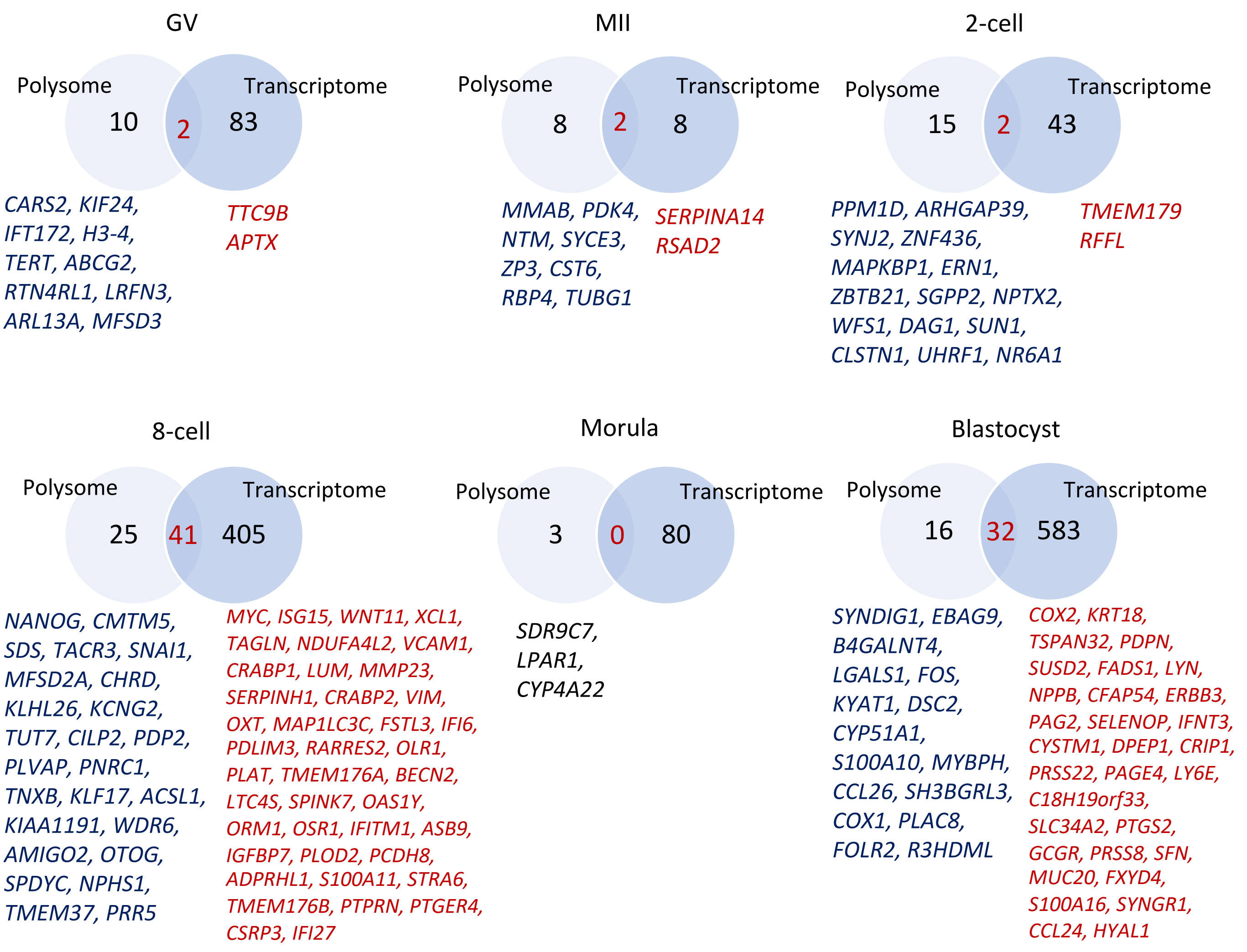
